# Genome editing human primary T cells with microfluidic vortex shedding & CRISPR Cas9

**DOI:** 10.1101/2020.02.26.960336

**Authors:** Justin A. Jarrell, Brandon J. Sytsma, Leah H. Wilson, Fong L. Pan, Katherine H.W.J. Lau, Giles T. S. Kirby, Adrian A. Lievano, Ryan S. Pawell

## Abstract

Microfluidic vortex shedding (*μVS*) can rapidly deliver mRNA to T cells with high yield. The mechanistic underpinning of *μVS* intracellular delivery remains undefined and *μVS*-Cas9 genome editing requires further studies. Herein, we evaluated a series of *μVS* devices containing splitter plates to attenuate vortex shedding and understand the contribution of computed force and frequency on efficiency and viability. We then selected a *μVS* design to knockout the expression of the endogenous T cell receptor in primary human T cells via delivery of CRISPR-Cas9 ribonucleoprotein (RNP) with and without brief exposure to an electric field (*eμVS*). *μVS* alone resulted in an equivalent yield of genome-edited T cells relative to electroporation with improved cell quality. A 1.8-fold increase in editing efficiency was demonstrated with *eμVS* with negligible impact on cell viability. Cumulatively, these results demonstrate the utility of *μVS* and *eμVS* for genome editing human primary T cells with Cas9 RNPs.

## Introduction

Intracellular delivery is a critical process in medicine and biology where exogenous materials are delivered across the cell membrane and into the cytosol. The intracellular delivery of constructs (i.e. DNA, RNA, protein and complexes) into cells allows for fundamental and exploratory research in cellular biology along with the synthesis of engineered cells as therapies^1^. The development of a simple and scalable microfluidic technology for intracellular delivery has the potential to dramatically improve the discovery process in research while also accelerating the development and manufacture of life-extending T cell immunotherapies like chimeric antigen receptor T cells (CAR-T)^2^ and T cell Receptor T cells (TCR-T)^3^.

*μVS* represents a simple and scalable approach to intracellular delivery^4^. Based on the fluid dynamics phenomena of vortex shedding, *μVS* is induced as fluid flows past a bluff body (i.e., a micron-scale post in a microfluidic device) creating alternating low-pressure regions downstream of the bluff body. The hydrodynamic conditions created by *μVS* are capable of permeabilizing the lipid bilayer of the cellular membrane to induce transient poration and permit intracellular delivery^4^. Despite these promising characteristics and potential, the fundamental physics contributing to *μVS* remains unclear. Thus, the development of *μVS* as an effective platform for cellular modification requires an improved understanding of vortex shedding to apply *μVS* to the intracellular delivery of additional constructs, cell types and conditions.

To this end, we developed a series of three-dimensional, transient, single-phase computational fluid dynamics models of *μVS*. We generated *μVS* designs that attenuate vortices through the addition of various ‘splitter plates’ between post columns of a six-column post array^5^. Using our computational fluid dynamics (CFD) model, we demonstrated that reductions in splitter plate lengths correlated with hydrodynamic conditions to enhance vortex shedding. These effects were subsequently validated through the fabrication of splitter plate devices and delivery of eGFP mRNA to recently activated primary human T cells via *μVS*. To explore its potential broader applications, we then applied *μVS* to the delivery of Cas9 and sgRNA targeting the T cell receptor alpha locus (TRAC-1), as a Cas9-RNP complex and successfully knocked out the expression of the endogenous T cell receptor of recently activated primary human T cells.

To better understand the influence of *μVS* on T cell phenotype, a direct comparison to electroporation was performed evaluating delivery efficiency, cell viability, proliferation, and levels of CD25, PD-1, and IFNg. This demonstrated that *μVS* results in an equivalent yield of genome-edited T cells shortly after intracellular delivery while maintaining the native T cell phenotype relative to electroporation. We also explored the addition of an electric field to *μVS* (*eμVS*) and increased initial editing efficiencies by 1.8-fold with negligible effect on T cell viability over the course of 14 days. Cumulatively, this demonstrates *μVS* and *eμVS* are robust alternatives to electroporation for genome editing recently activated human primary T cells.

## Results

### Hydrodynamic simulations demonstrate reductions in vortex shedding with the inclusion of splitter plates

Splitter plates of various length were incorporated into a previously reported *μVS* device design^4^ to evaluate the mechanistic underpinning of microfluidic vortex shedding. Empirical measurements of the inlet and outlet pressures and single-phase fluid properties indicated an object Reynolds number of 127 similar to the previously reported experiments and two-dimensional simulations^4^. Splitter plate ratio was defined as the ratio between the length of the splitter plate (i.e. a thin object between two consecutive columns of posts) and the center-to-center distance of two consecutive cylindrical posts in the flow-wise direction. Separation ratio was defined as the ratio of the distance between the center of the cylindrical post (in the 2D-plane, this is the center of the circle) and the leading edge of the splitter plate to the diameter of the cylindrical post^5^. Together, these ratios resolved five device designs, each with a unique splitter plate ratio: 0, 0.25, 0.50, 0.75, and 1.0 (Supplemental Figure 1). The respective separation ratios for these devices are found in Table 1.

**Table 1.** Summary of splitter plate device designs and simulations

| Device Designs |  |  | Simulations |  |  |
| --- | --- | --- | --- | --- | --- |
| Design | Splitter Ratio | Separation Ratio | Vortex Frequency (kHz) | Spanwise Fluctuating Hydrodynamics Forces ($\mu N$) | Post-Near Wake Indicator (PNWI) |
| SIM01 | 1.0 | 0 | -- | 3.9 | 1% |
| SIM02 | 0.75 | 0.75 | -- | 6.4 | 2% |
| SIM03 | 0.50 | 2 | -- | 8.7 | 3% |
| SIM04 | 0.25 | 3.25 | 36.0 | 134 | 40% |
| SIM05 | 0 | 10 | 13.5 | 336 | 100% |

CFD simulation results provided computed flow data including hydrodynamic pressure, flow velocity, and microfluidic vorticity to evaluate vortex strength, structures and shedding behavior of simulated flows within *μVS* devices (Supplemental Method 1). Simulations indicated that both spanwise hydrodynamics fluctuating force and Post Near Wake Indicator (PNWI, Supplemental Method 2) increased with reductions in splitter plate ratios (Table 1, Supplemental Figure 2A & B). In fact, devices with a splitter plate ratio ≥0.50 showed minimal to no vortex shedding due to computed vortex attenuation. The total spanwise hydrodynamic fluctuating forces in 1.0, 0.75, 0.50 splitter plate ratio devices was determined to be approximately 3.9 *μ*N, 6.4 *μ*N, and 8.7 *μ*N, respectively or order(s) of magnitude lower than that of the shorter or absent splitter plates. The magnitude of these fluctuations were much greater with 0.25 and 0.0 splitter plate ratio devices at approximately 134 *μ*N and 336 *μ*N, respectively. Similarly, the PNWI values of 0.25 and 0.0 splitter plate ratio devices were also determined to be greater than devices with splitter plate ratios ≥0.50 at 40% and 100%, respectively (Table 1). Both increased PNWI and fluctuating force results indicate a substantial increase in vortex ‘strength’ with a decrease in splitter plate lengths. Based on this, we observed the highest vortex ‘strength’ in simulated devices without splitter plates. The measurements also suggested that when splitter plates were placed sufficiently close to the trailing edge of the cylindrical posts, vortex shedding failed to develop. In general, our simulations demonstrated a consistent inverse correlation in PNWI, spanwise hydrodynamics fluctuating forces, and the splitter plate or separation ratio.

In addition, the simulation results also provided evidence of vortex structures present in the device as quantified by the vortex shedding frequency. Vortex shedding frequency was quantified from the periodic form of oscillation of the vortex structures behind the posts. It was found that only 0.25 and 0.0 splitter plate ratio devices exhibited consistent periodic oscillation of the dominant vortex shedding, at rates of 36 kHz and 13.5 kHz, respectively (Table 1, Supplemental Figure 3). Minimal to no vortex structures were observed in the 0.5, 0.75, or 1.0 splitter plate ratio configurations with zero vortex shedding frequencies detected. It should be noted that these simulations are for single-phase flow that may only be representative of the flow of suspended cells through a *μVS* device. Furthermore, vortex shedding spectral analysis also suggested that vortex suppression occured when the separation ratio was less than 3.25 (SIM01-03, splitter plate ratio 1.0 - 0.5), similar to previous reports^5^. Numerical analysis also revealed that vortex formation inside the wake region occurred as a result of two (‘pair’) of counter-rotating vortices developed behind a post (Supplemental Figure 4). In the 0.0 splitter plate ratio device, the wavelength of its vortex pairs was approximately 6 times the post diameter (240*μm*). This is consistent with the computed vortex convection speed of 3 m s^-1^ at the frequency rate of 13.5kHz based on wavelength-frequency relationship. However, in the 0.25 splitter plate ratio device, the presence of this splitter plate (with a separation ratio of 3) interfered with oscillation downstream of the posts while full oscillating wavelengths were undetectable in the 1.0, 0.75, and 0.5 splitter plate ratio devices (Supplemental Figure 2B & 5). Devices simulated with a separation ratio of 3 (SIM04, 0.25 splitter plate ratio) resulted in a higher frequency of vortex oscillation with lower spanwise hydrodynamic forces when compared to SIM05 (0 splitter plate ratio) (Table 1, Supplemental Figure 3). Overall, the inclusion of various splitter plates allowed for the incremental attenuation of vortex shedding in a manner that enabled us to computationally determine vortex shedding frequency and total spanwise fluctuating hydrodynamics forces. Subsequently, we were able to compare these computed results to biological experiments in order to assess correlations between cell viability, mRNA delivery efficiency, vortex shedding frequency and/or total spanwise fluctuating hydrodynamics forces. Achieving a similar result empirically through high speed imaging and/or micro-particle image velocimetry is simply not feasible due to the sampling frequency and cell media composition requirements, respectively.

The hydrodynamic pressure distribution contours indicated that the fluid pressure remains uniform in regions between consecutive post columns. Such hydrodynamic pressure distribution was unperturbed in the presence of vortex shedding structures in 0.0 and 0.5 splitter plate ratio devices (Supplemental Figure 1). Pressure uniformity was self-sustained by a pair of counter-rotating vortices with similar strengths oscillating at both symmetrical sides of the post (Supplemental Figures 1 & 2). The pressure distribution remaining undisturbed in the presence of vortex shedding inferred that the flow dynamics in the device would remain unchanged. Therefore, it is speculated that the flow rates for all splitter plate ratio devices remained consistent at a given applied pressure.

Vortex structure behavior was also detected by measurement of Q-criterion iso-surfaces (Supplemental Figure 6). Q-criterion analysis indicated that the vortex structures were oscillating at higher amplitude in the downstream post columns (4-6) compared to upstream columns (1-3) in the flow cells of the 0.0 and 0.25 splitter plate ratio devices. Based on this, it is speculated that the number of post columns included in a device design will impact the overall device performance (i.e. delivery efficiency) due to vortex amplitude enhancement. Furthermore, the self-sustained vortex fluctuations behavior in the downstream columns are also thought to further enhance device performance as an intracellular delivery method.

### Vortex shedding correlates with delivery efficiency & cell viability

Our CFD simulations highlighted key hydrodynamic features that enhance vortex shedding and could promote intracellular delivery. Therefore, we evaluated these results empirically by fabricating and testing *μVS* devices with splitter plates of various lengths for intracellular delivery of eGFP mRNA to recently activated primary human CD3+ T cells (Supplemental Figure 1). As predicted in our simulations, we observed incremental decreases in eGFP expression with increases in splitter plate lengths, ranging from 51.8% to 11.8% expression between 0 and 1.0 splitter plate ratio devices, respectively, (Figure 1A, Table 2). Conversely, we observed an increase in cell viability with decreases in splitter plate lengths attributed to the attenuation of fluid forces that diminish vortex shedding. However, unlike eGFP expression, the influence of vortex shedding on cell viability were stepwise, or non-incremental, with the largest increases in viability occurring between 0.5 and 0.75 splitter plate ratio devices with 44.6% and 66.9% live cells quantified, respectively (Figure 1B, Table 2). When compared to our numerical CFD simulations, our experimental results indicate that the presence and strength of computed total spanwise fluctuating hydrodynamic forces is positively correlated with intracellular delivery efficiency with a stepwise decrease in initial cell viability at approximately the onset of vortex shedding.

**Figure 1.**
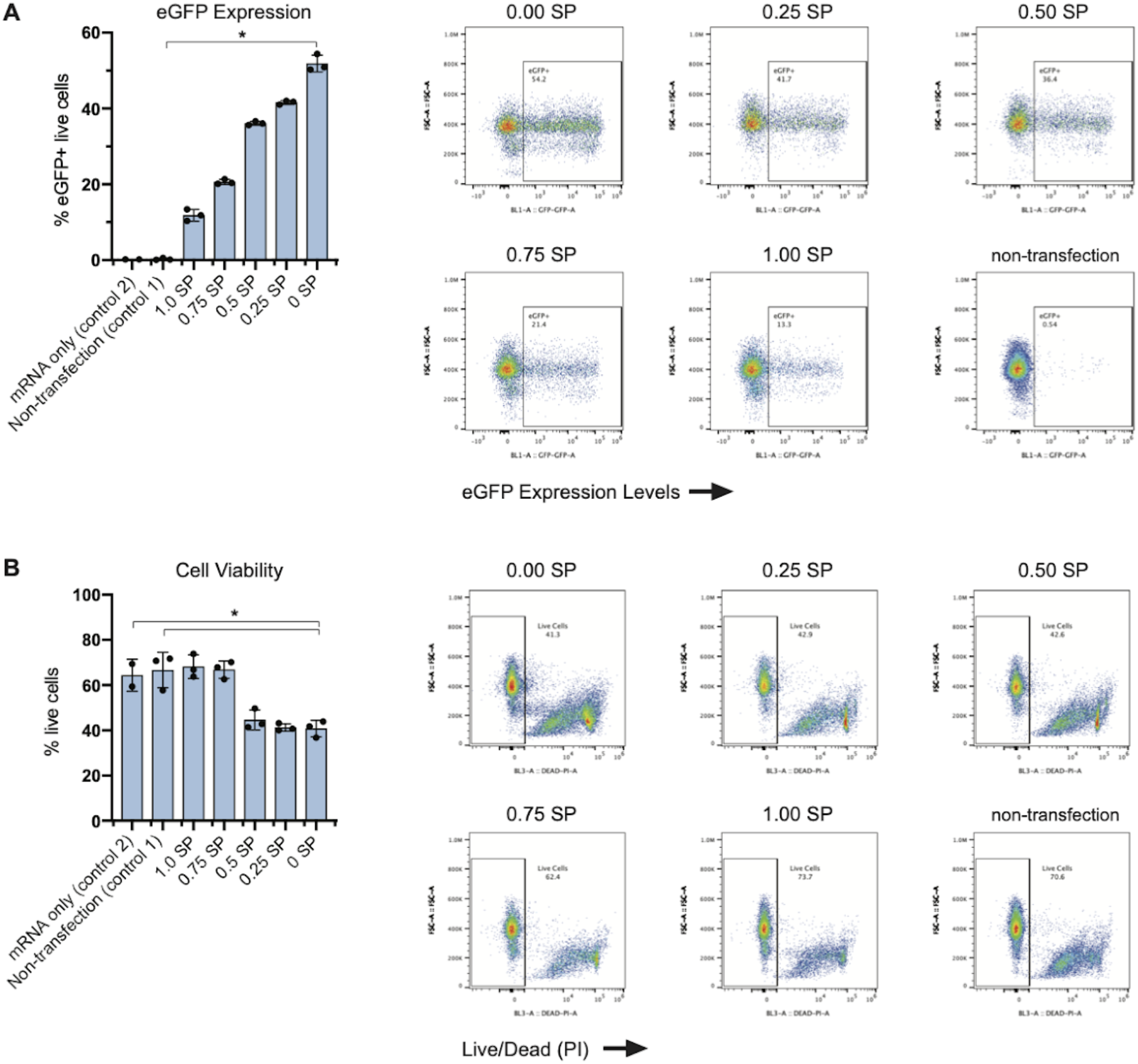
Magnitude of vortex shedding correlates with eGFP expression and cell viability. eGFP-encoding mRNA was delivered to activated primary human CD3+ T cells via *μVS* with microfluidic devices containing splitter plates of various lengths. SP ratio is defined as the ratio between the length of the splitter plate and the center-to-center distance of two consecutive cylindrical posts in the flow-wise direction. Levels of (A) eGFP expression and (B) cellular viability were quantified at 24 hours post-transfection. Data represent the mean ± standard deviation of n = 3 samples per condition. Dot plots representative of triplicate samples with SP ratios and control indicated in each plot. *P < 0.05 by Kruskal-Wallis.

**Table 2.** eGFP expression and cell viability at 24 hours post-*μVS* with splitter plate devices

| Device Designs |  |  | Expression & Viability |  |
| --- | --- | --- | --- | --- |
| Design | Splitter Ratio | Separation Ratio | Efficiency (%) | Viability (%) |
| SIM01 | 1.0 | 0 | 11.8 ± 1.55 | 68.2 ± 5.19 |
| SIM02 | 0.75 | 0.75 | 20.6 ± 0.68 | 66.9 ± 3.85 |
| SIM03 | 0.50 | 2 | 36.1 ± 0.50 | 44.6 ± 4.44 |
| SIM04 | 0.25 | 3.25 | 41.6 ± 0.53 | 41.3 ± 1.62 |
| SIM05 | 0 | 10 | 51.9 ± 2.21 | 40.8 ± 3.63 |

### Genome editing with Cas9 & *μVS*

To explore the broader application of *μVS* beyond intracellular delivery of eGFP mRNA, we evaluated the utility of *μVS* for CRISPR-based T cell engineering relative to electroporation. Using both *μVS* and electroporation, we delivered Cas9 and single-guide RNAs (sgRNAs), as a Cas9-RNP complex targeting the first exon of the TCR-alpha constant region (TRAC-1) to 5.0 x 10^6^ activated primary human CD3+ T cells. As a proxy for delivery efficiency, we evaluated TCRα/β and CD3 co-expression in the live cell populations. We quantified co-expression at 4 days after transfection and observed a significant decrease in TCR/CD3 expression with *μVS* and electroporation, with an average editing efficiency of 25% and 95% of live T cells, respectively, compared to cells transfected with a non-target sgRNA-Cas9 RNP and non-transfection controls (Figure 2A). These reductions in TCR/CD3 expression remained stable at days 7, 10 and 14 post-transfection, with less than 2% variability in TRAC-1 editing efficiencies for both methods (Figure 2A). A statistically significant difference in the percentage of TCR/CD3-T cells was observed between *μVS* and electroporation for all time points post-transfection.

**Figure 2.**
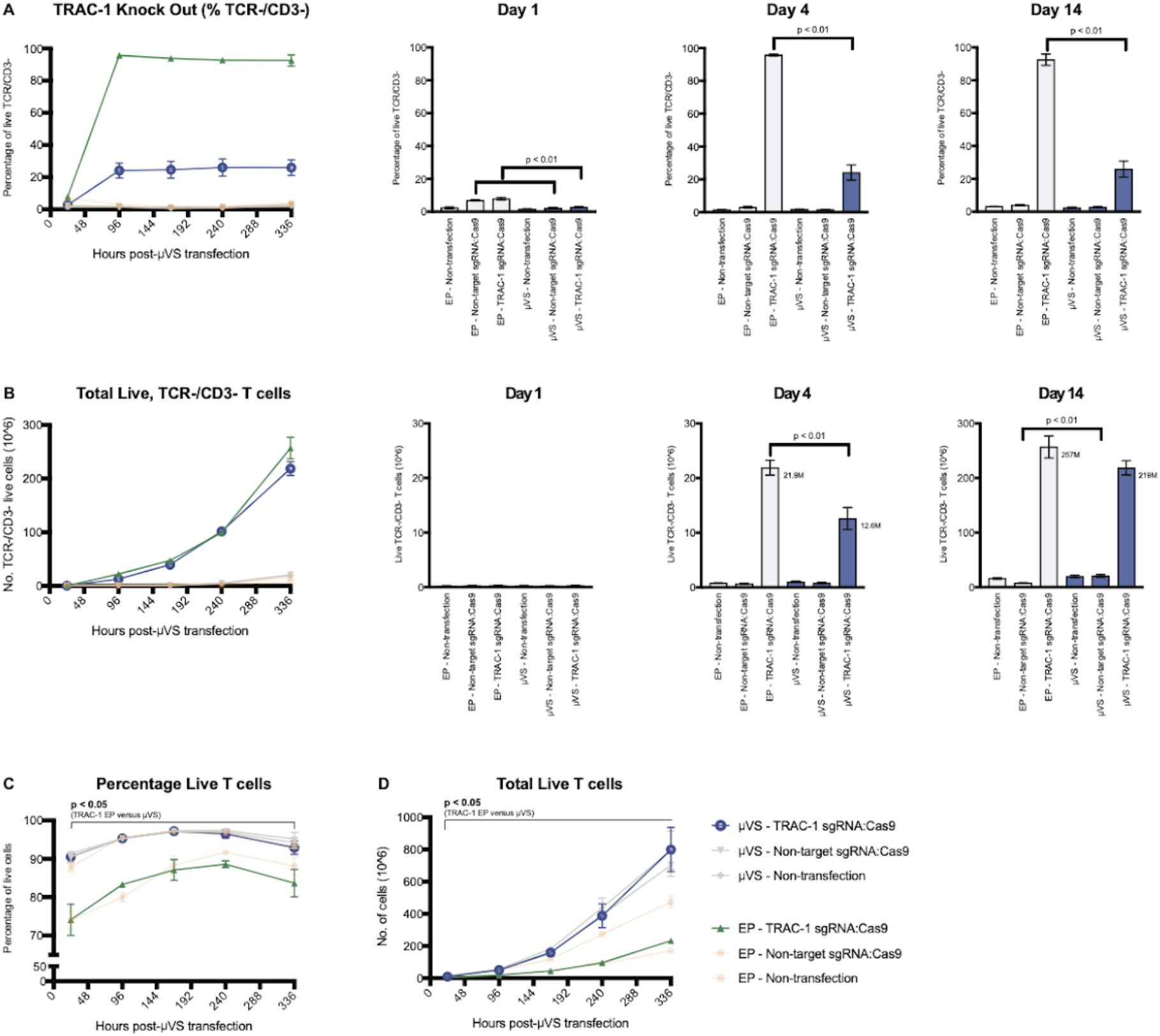
Persistent knockout of endogenous TCR in primary human T cells with CRISPR and *μVS*. Cas9 and locus-specific TRAC-1 gRNAs (as Cas9-RNP complexes) were delivered to activated primary human CD3+ T cells via *μVS* or electroporation. Quantification of **A**, percentage of TCR knock-out cells, **B**, total viable, TCR knock-out cells, **C**, percentage of viable T cells and **D**, total viable T cells were performed at Days 1, 4, 7, 10 and 14 post-transfection. TCR expression was measured via CD3 and TCRa/b co-staining. TCR knockout was quantified as a percentage of CD3- and TCRa/b-double negative cells and the total number of live cells. Propidium iodide exclusion gating and event collection rate were used to measure viability and cell concentration, respectively.. Non-targeting gRNA as a Cas9-RNP complex served as a non-editing control. Non-transfected samples served as a negative control. Data represents mean ± SD of biological triplicate. P-values by unpaired, two-tailed, heteroscedastic T-tests.

In addition to quantifying TCR editing efficiency, we also evaluated the number of TCR knockout cells and the effects on cell viability over time for each transfection method. While we observed a statistically significant difference in TCR editing efficiency between *μVS* and electroporation, both methods yielded equivalent numbers of modified T cells by day 7, with these levels maintained at days 10 and 14 post-transfection. At 24 hours post-transfection, we detected >90% viable cells in samples processed via *μVS* compared to 73% with electroporation. Peak viabilities of 98% and 85% were observed at days 7 and 10 with *μVS* and electroporation, respectively. Unlike electroporation, cells processed via *μVS* had viabilities comparable to non-transfection control samples for the duration of the experiment (Figure 2B). In addition, we also observed a consistent enhancement in cell viability with *μVS*, with a 9-16% increase in the percentage of live cells observed with *μVS* compared to electroporation at each time point post-transfection (Figure 2C). Furthermore, we quantified the number of live T cells after transfection and observed a 3.4X increase in live T cells achieved with *μVS* (8 x 10^8^ cells) compared to electroporation (2.3 x 10^8^ cells) at day 14 post-transfection. Similar to the non-transfection controls, *μVS*-processed samples experienced a 160-fold expansion in the number of live T cells at day 14 compared to the numbers observed at 24-hours post-transfection. Unlike *μVS*, electroporation led to a substantial reduction in the proliferation rates of live T cells observed after transfection, with a 46-fold expansion in live T cells quantified at day 14 and proliferation curves that did not track with corresponding non-transfection controls (Figure 2D).

Electroporation has been shown to augment the expression of functional T cell surface markers and cytokines associated with their diminished function in vitro^6^. Therefore, to better understand the effect of transfection methods on T cell phenotypes, we quantified the expression of CD25 and PD-1 in cells transfected via *μVS* compared to electroporation. Statistically significant differences in CD25 expression levels were observed on day 4, 10 and 14 post-transfection when delivering TRAC-1 Cas9-RNP complexes to T cells via *μVS* and electroporation. We observed a consistent downregulation of CD25 expression ranging from 5-10% in live T cells processed via electroporation compared to *μVS* as well as non-transfection controls in early time points. Conversely, a statistically significant elevation in the levels of CD25 expression from 5-15% of live T cells was also observed in electroporated samples compared to *μVS* and non-transfection controls at days 10 and 14 post-transfection. A similar trend was also observed in TRAC-1 KO live cell populations. However, unlike electroporation, no significant difference in CD25 expression was detected in cells transfected with TRAC-1 or non-targeting RNPs via *μVS* compared to non-transfected control cells, indicating that the expression of CD25, an important component of the IL-2 receptor and early T cell activation, is minimally impacted by *μVS* (Figure 3A).

**Figure 3.**
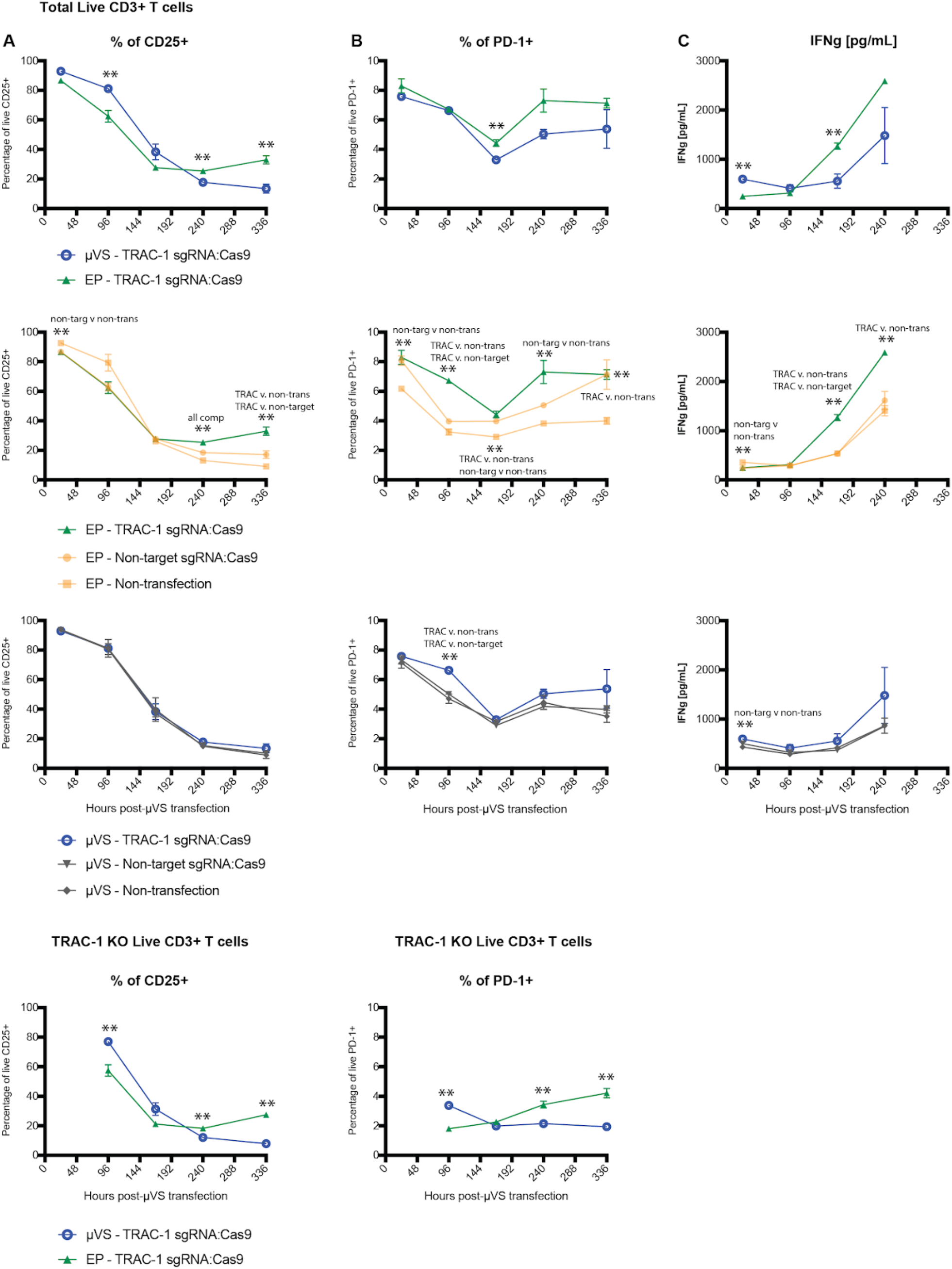
Surface marker expression and cytokine secretion levels minimally perturbed with CRISPR and *μVS*. **A**, CD25, and **B,** PD-1 expression levels were quantified in cells transfected with *μVS* or electroporation. Surface marker expression levels were measured in the total live cell and TRAC-1 KO live population via flow cytometry at Days 1, 4, 7, 10 and 14 post-transfection. **C**, ELISA quantification of IFNg supernatant levels in transfected and non-transfected cells at Day 1, 4, 7, and 10 post-transfection. Non-targeting gRNA as a Cas9-RNP complex served as a non-editing control. Cells mixed with P3 buffer or Opti-MEM buffer media that were not transfected served as non-transfection controls for electroporation and *μVS* samples, respectively. Data represent mean ± SD of 3 wells. P-values by unpaired, two-tailed, heteroscedastic T-tests.

Similar to CD25, the expression of PD-1, a T cell marker of early activation and exhaustion, is also significantly affected by electroporation, with elevated levels observed at days 7 and 10 post-transfection compared to *μVS*. Similar to total live T cells, a statistically significant increase in PD-1 expression was also observed at the same time points in TRAC-1 KO cells generated vial electroporation compared to μVS. Furthermore, increased levels of PD-1 expression were observed in cells transfected with TRAC-1 or non-targeting RNPs via electroporation compared to non-transfection controls at days 4, 7, and 14 post-transfection. In contrast, PD-1 expression levels in live T cells were minimally perturbed when transfected via *μVS*, with PD-1 expression comparable to levels detected in T cells of non-transfection control samples (Figure 3B). These results highlight that the process of electroporation augments PD-1 expression and potentially promotes exhaustion of T cells, while *μVS* preserves expression levels most similarly to the unperturbed native T cell state.

In addition to surface marker expression, we also quantified the supernatant levels of pro-inflammatory cytokine, interferon-gamma (IFNg), for 14 days following *μVS* and electroporation. Like PD-1 and CD25 expression, we observed a statistically significant increase in the concentration of IFNg present at 7 days post-transfection with electroporation compared to *μVS*. A similar trend was also observed in our comparison of electroporated versus non-transfection samples, where a statistically significant increase in the levels of IFNg were detected at days 7 and 10. By day 14 post-transfection, IFNg levels were 2.3-fold higher in electroporated samples compared to *μVS*-processed samples when delivering TRAC-1 targeting RNP, and 1.5-fold higher when delivering non-targeting RNP (Figure 3C). These elevations in IFNg production in the absence of TCR engagement highlight an additional change in T cell phenotype that could alter cell function as a result of electroporation, with a preservation of the normal levels of IFNg maintained with *μVS*.

### Efficient genome editing with CRISPR Cas9 & *eμVS*

Our experiments with *μVS* resulted in an equivalent yield of CRISPR-edited T cells compared to electroporation. Therefore, we hypothesized that incremental enhancements in knockout efficiencies with the maintenance of high cell viabilities could result in an increased number of edited T cells surpassing electroporation. Thus, we sought to increase Cas9 RNP uptake in T cells via integration of interdigitated electrodes in our microfluidic devices to apply an electric field alongside *μVS*, called *eμVS* (Figure 4).^7,8^ Preliminary experiments with OptiMEM as a processing media indicated a preferred applied electric field strength of 2.25 kV cm^-1^ where significant bubbling and clogging was observed above 2.25 kV cm^-1^ (data not shown). *eμVS* resulted in a 1.8-fold increase in editing efficiency (38.4%) relative to *μVS* (21.4%) while maintaining greater than 80% cell viability 24 hours after processing (Figure 5A). Interestingly, there was no statistically-significant difference in cell viabilities for *μVS*, *eμVS* and the non-transfection control within 72 hours after processing with greater than 90% viable cells observed through day 10 for all conditions (Figure 5B).

**Figure 4.**
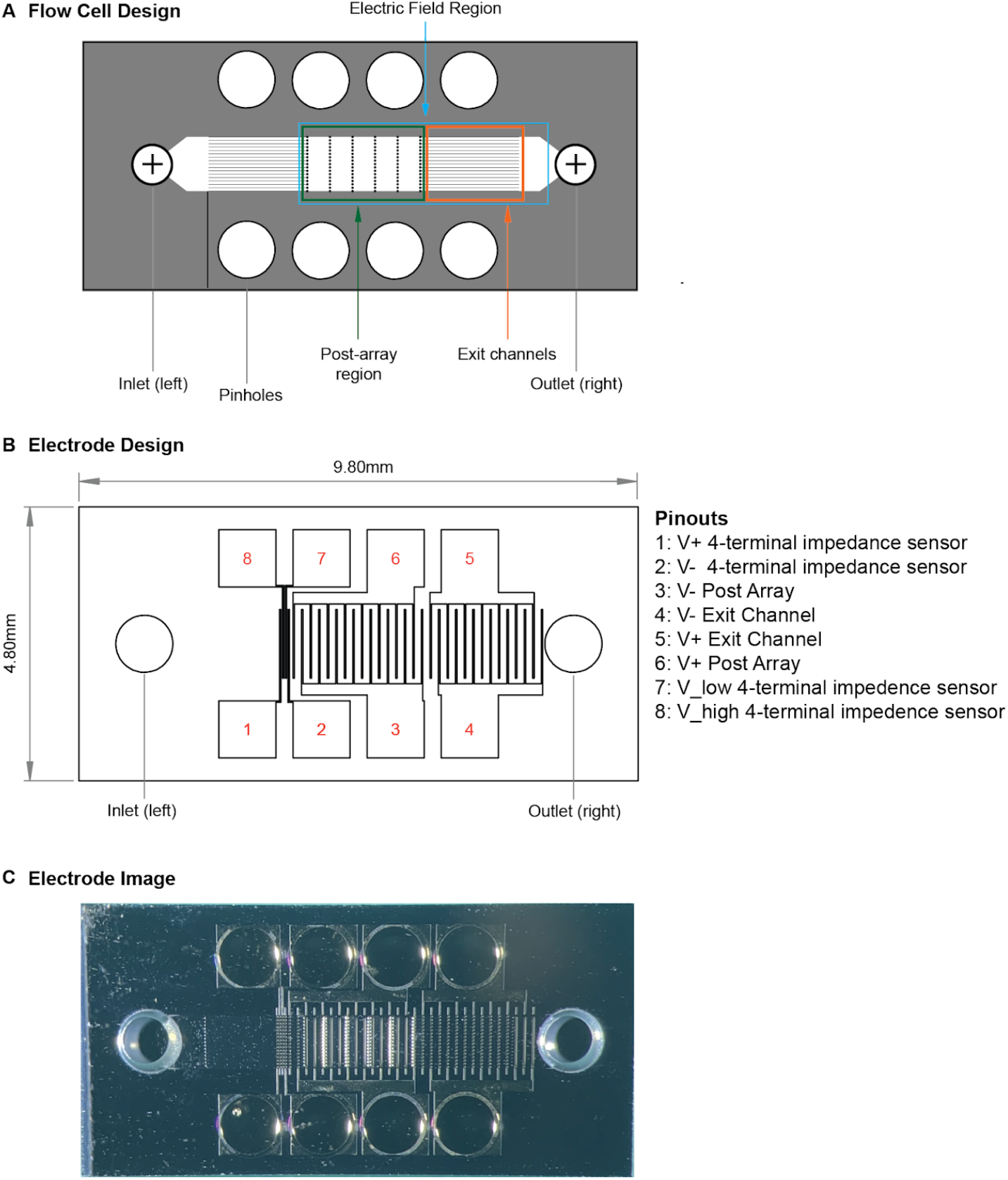
e*μVS* device and electrode designs. **A**, 4.8 x 9.8 mm deep reactive ion etched silicon substrate contained a previously reported^4^ 40 μm deep flow cell and containing an array of 40 μm diameter posts without splitter plates (see Supplemental Figure 1B) and 1 mm pinholes for electrical access. **B**, each flow cell substrate is anodically bonded to a laser-machined borofloat lid. The laser-machined lid contained an electrode array for (1) measuring the electric field with a 4-terminal impedance sensor, (2) an interdigitated electrode array over the post array and (3) an interdigitated electrode array over the exit channels. Both interdigitated electrode arrays contained 25 μm wide electrodes spaced 125 μm apart. When (2) and (3) were operated at the same time with a 30V DC offset, this enabled an up to 2.4 kV cm^-1^ applied electric field to pulse the cells 29 times over a total distance of 4.35 mm. **C**, Optical micrograph of a 4.8 x 9.8 mm anodically-bonded *eμVS* device taken from the optically-transparent borofloat lid side of the device containing the 800 μm laser machined fluidic inlet and outlet.

**Figure 5.**
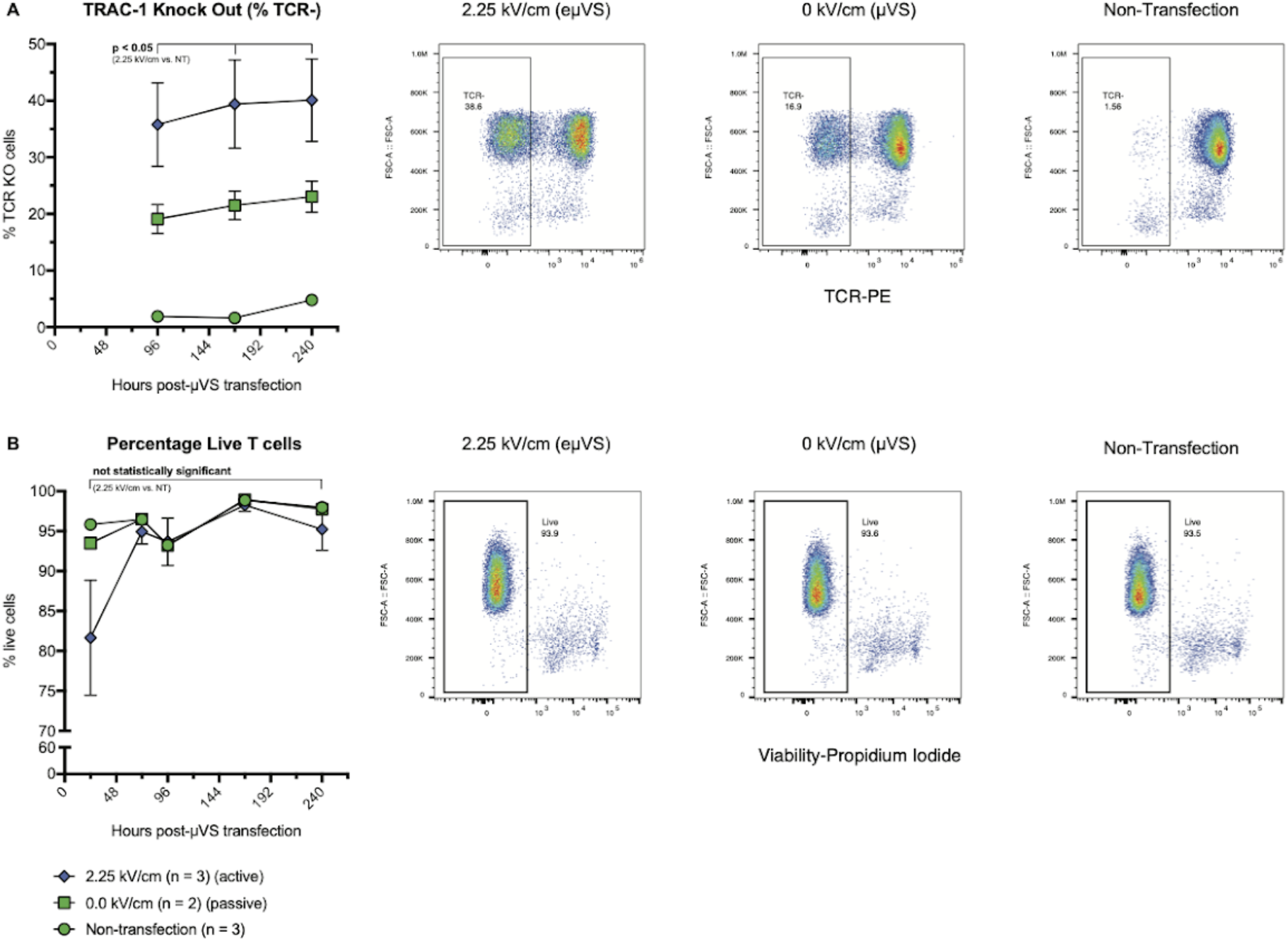
Application of electric field with *μVS* (*eμVS*) promotes intracellular delivery and TRAC-1 knockout. TRAC-1-targeting Cas9-RNPs were delivered to activated primary human CD3+ T cells via *eμVS* with an applied electrical field. Quantification of **A**, percentages of TCR knock-out cells and **B**, viable T cells were quantified via flow cytometry at multiple timepoints post-transfection. TCR knockout levels were measured via TCRa/b staining and quantified as a percentage of TCRa/b-negative cells and the total number of live cells. *eμVS* with no applied electric field (0 kV cm-1) and non-transfection samples served as controls. Data represent the means ± SD of n ≥ 2 samples per condition. Statistical analysis performed between groups indicated on graphs. Dot plots representative of replicate samples. P<0.05 by unpaired, two-tailed, heteroscedastic T-tests.

## Discussion

### Splitter plate simulations & experimentation

The use of splitter plates of variable lengths allowed us to analyze the influence of vortex shedding on delivery efficiency and cell viability in a manner where the influence of pressure changes and vorticity experienced by cells could be reasonably decoupled. Splitter plate designs were able to completely attenuate vortex shedding with a splitter ratio less than 0.5 with only partial levels of vortex shedding occuring in devices with a splitter ratio of 0.25. The stepwise reduction in cell viability observed at the 0.5 splitter ratio indicates single-phase simulation may not account for the onset of vortex shedding with particle-laden flows or those containing suspended cells. The highest levels of vortex shedding was observed in a previously reported *μVS* device design with no splitter plate^4^. All of this is in agreement with the fundamental splitter plate studies performed by Unal and Rockwell at similar object Reynolds numbers.^5^

Interestingly, a trend between these simulation parameters (i.e. total spanwise hydrodynamics fluctuation force and the PNWI) and the eGFP expression efficiency of mRNA and T cell viability is observed. As the splitter ratio of the *μVS* device decreases, the delivery efficiency increases, which is correlated to the rise of the total spanwise hydrodynamics fluctuation force and vortex shedding frequency in simulations. Similarly, previous studies have also demonstrated a negative correlation between delivery efficiency and cell viability^4^. In the *μVS* devices containing splitter plates, a comparable trend emerged with enhanced delivery efficiency correlating with a reduction in splitter plate lengths and reduced 24 hr cell viability.

Discontinuities were also observed in simulation and empirical results highlighted in our analysis of the *μVS* device with a 0.5 splitter ratio. Despite simulating an absence of vortex shedding and minimal spanwise fluctuating hydrodynamic forces, eGFP expression was observed in the 0.5 splitter plate ratio devices. We speculated that the delivery of eGFP mRNA in cells could be due to the presence of early-onset, cell-induced vortex shedding that was not captured in our single-phase simulations in addition to the pressure changes caused by constricted flow between posts. Single-phase simulations are limited in their ability to capture nuanced vortex shedding in the presence of suspended cells and will likely require transient, multi-phase approaches to simulate suspensioned cells to better understand the contribution of vortex shedding on delivery efficiency. Multi-phase simulations with particle-laden flows are currently under development and further development of high-speed imaging at or above the Nyquist sampling frequency is required to verify those simulations.

A key limitation in our current study also includes not directly correlating computational and numerical parameters to empirical results on biological systems (i.e. cell age and growth, activation time etc.). Improvements to this framework would include the empirical quantification of transient hydrodynamic forces experienced by suspended cells for each *μVS* device design using multi-phase simulation studies of particle-laden flows. Despite these limitations, our study serves as a foundation to correlate CFD simulations and biological experiments to better understand *μVS* as a novel method for intracellular delivery.

### Genome editing with CRISPR-Cas9 & *μVS*

The CRISPR-Cas9 genome editing system has emerged as a promising approach to generate cell-based immunotherapies without the use of viral vectors along with an array of other practical applications.^9^ Recently, T cells engineered to express an exogenous T cell receptor targeting tumor antigens via delivery of Cas9-RNP and DNA templates have been shown to mount effective anti-tumor responses *in vitro*, in murine models^10^, and in early stage clinical trials^3^. To explore the utility of *μVS* as an intracellular delivery method for CRISPR-Cas9 genome editing, we delivered Cas9 protein complexed to sgRNA targeting the T cell receptor alpha locus (TRAC-1) to knock out the expression of the endogenous T cell receptor and permanently modify recently activated primary human T cells. To demonstrate the feasibility of *μVS* as a tool for recently activated T cell genome editing, we performed a head-to-head comparison of *μVS* with an established electroporation protocol^10^. With *μVS,* we demonstrated ~25% editing efficiency in live T cells at 4 days post-*μVS*, compared to 95% observed in electroporated samples. Both transfection techniques resulted in stable genome editing which persisted for an additional 10 days for the duration of the experiment. Cell viability was also minimally impacted with *μVS*, with >90% viable cells achieved at 24 hours and maintained for all time points thereafter, exceeding the product release criteria for the clinical use of current CAR-T cell therapies^2^. Unlike *μVS*, a substantial decrease in cell viability was observed as a result of electroporation, with ~75% viable cells observed at 24 hours post-transfection and a 9-16% decrease in viable cells observed at all time points compared to *μVS* and control samples. These significant and permanent reductions in cell viability are hallmarks of electroporation, and have already proven to be a significant challenge in its adoption as a practical intracellular delivery method for cell therapy manufacturing.^6,11^

Accompanying delivery efficiency and viability as important factors to generate effective cell therapies, the ability to synthesize a sufficient number of modified cells to dose patients is a critical factor in successful treatment in the appropriate time frame^2,3^. Because of this, the expansion rate of genome edited T cells following transfection is of critical importance. While we observed a nearly 4-fold increase in TRAC-1 editing with electroporation compared to *μVS*. the superior rate of expansion in *μVS* samples produced an equivalent number of genome edited T cells by 7 days after transfection. In addition, a 1.8-fold increase in TRAC-1 editing efficiency was achieved with the inclusion of an electric field coupled with *μVS*, *eμVS.* indicating that an enhancement in genome editing can be achieved with minimal impact to cell viability. Interestingly, TRAC-1 editing efficiencies demonstrated with *eμVS* were comparable to those reported in the first-in-human phase I clinical trial with multiplex CRISPR-Cas9 editing^3^. These results highlight that *μVS* and *eμVS* can likely be leveraged to minimize the time required to generate sufficient numbers of edited T cells for clinical applications beyond electroporation, without compromising cell quality. This is particularly important in light of a recent *in vivo* study demonstrating that a reduction in cell culture time (i.e. 3 days) enhanced anti-leukemic activity in CAR-T cells at a 6-fold lower dose^2^. The rapid gene editing time frames coupled with the development of *μVS* design with higher cell processing capacity, highlights the potential of *μVS* and *eμVS* to substantially reduce manufacturing times required to generate T cell immunotherapies, like CAR-T and TCR-T therapies at clinical and commercial scales.

### Enhanced genome editing with CRISPR Cas9 & *eμVS*

The challenge of intracellular delivery stems from the impermeability of the plasma membrane. Electroporation seeks to overcome this challenge by creating temporary pores in the cell membrane through the application of a short, high voltage electric pulse (i.e. 100 μs, 5 kV cm^-1^)^12^. As the phospholipid bilayer is charged by the electric field, the transmembrane potential will reach a certain threshold where pores begin to form on the cell surface^13^. The size and duration of these transient pores are largely dependent on the strength and pulse width of the electric field. Studies show that the larger the pore is the more rapidly it will close once the electric field is removed^13^. To take advantage of this brief window of large-molecule permeability, electroporation (via Lonza’s Nucleofection) uses a second, longer pulse with an assumed lower electric field strength (4 A*cm^-2^ and 40 ms) to augment the uptake of extracellular cargo^12^. This second longer pulse provides electrophoretic force which acts on the charged cargo in suspension to drive it through the transient pores and into the cytoplasm. *μVS* uses hydrodynamic forces over an estimated 200 μs period, rather than a 100 μs, 5 kV cm^-1^ electrical pulse, to permeabilize the plasma membrane. The temporary pores are formed, however, *μVS* must rely on diffusion to move cargo into the cell. Here, we demonstrate the utility of *eμVS*. the combination of the hydrodynamic conditions of *μVS* along with a specifically tailored electrical field, in delivering Cas9-RNP to recently activated primary human T cells. Assuming an average cell speed of 10 m s^-1^, the *eμVS* device in this study would result in an estimated total electric field exposure time of 435 μs or 644-fold less than with continuous flow microfluidic electroporation^14^. Relative to electroporation, *eμVS* replaces the 100 μs, 5 kV cm^-1^ electric field with gentle hydrodynamic membrane poration step^4^ and enhances uptake with an electrophoretic field over a 92-fold reduced total electric field exposure time^12^. Cumulatively, this enables *eμVS* to efficiently deliver CRISPR Cas9 genome editing constructs to T cells while also minimizing impacts on cell viability relative to various forms of electroporation.

Importantly and during preliminary studies, it was determined that commercial electroporation buffers (such as those used by the Lonza Nucleofector and BTX Electroporation system) are extremely harsh on T cells after exposure times exceeding 30 minutes. For *μVS* and *eμVS*, however, we relied on commercially-available OptiMEM that does not significantly impact T cells viabilities with longer incubations and is available as a GMP-grade reagent. This may further enable *μVS* and *eμVS* as a practical and gentle intracellular delivery platform. Additional work is required to determine the influence of different processing media and their respective conductivities on *eμVS* performance.

Another important consideration during *eμVS* is the electric field. When using this *eμVS* device and the above protocol, cells are pulsed with 29 electric fields about 125 μm in length with a 25 μm gap between fields due to the electrode array geometries (see Figure 4B,C). This is similar to exposing cells suspended in OptiMEM to a 2.25 kV cm^-1^ applied electric field at 66.7 kHz with an 83.3% duty cycle for 29 pulse widths over 435 μs. In practice, the use of a 4-terminal impedance sensor (Figure 4B) could be used to measure the electric field strength. Varied flow speeds and cell trajectories as a result of vortex shedding in the post array region are likely to create a varied electric field exposure and this may result in the greater error bars seen with *eμVS* relative to *μVS* (Figure 5A). Reducing cell-to-cell processing variability for *eμVS* could be achieved by determining the minimum flow cell width that allows for efficient *μVS*. This would be similar to the above splitter plate study, but with reduced flow cell widths rather than splitter plates increasing in length. This would ensure less variability in cell trajectories due to recirculation resulting in more uniform electric field exposure. This minimal flow cell width could then be used for applications requiring low numbers of genome edited cells in discovery-stage workflows like high throughput screening and this same flow cell geometry could then be arrayed across a substrate for applications where significant numbers of genome edited cells are required like cell therapy^3^.

## Conclusion

Our investigation of microfluidic vortex shedding via simulations and experimentation provides a better understanding of the fluid dynamics contributing to *μVS* as a method for intracellular delivery. We demonstrated a novel application of *μVS* for genome editing activated human T cells with the CRISPR-Cas9 genome-editing system. Compared to electroporation, *μVS* resulted in superior cell viability and proliferation, and preservation in cell surface marker expression and cytokine secretion levels relative to non-transfection cell controls. Despite discrepancies in editing efficiency, we observed an equivalent number of genome edited T cells with *μVS* and electroporation. Further, we demonstrated enhanced CRISPR-based gene editing through e*μVS*, the application of *μVS* coupled to a brief electric field, with a 1.8-fold increase in TCR knockout editing efficiency with low impact on cell viability.

Studies for transgene insertion using (1) Cas9-RNPs with different DNA templates and targets^10^ and (2) a transposon/ase system^15^ with *eμVS* are currently ongoing. Future efforts to further appreciate the mechanistic underpinnings of *μVS* will focus on direct comparisons of multi-phase, particle-laden CFD simulations with high-speed imaging and will be used to better study the effects of fluctuating lift force and PWNI on intracellular delivery. These results will expedite the development of *μVS* and *eμVS* devices optimized to deliver a variety of constructs and cell types. Additional studies into the transcriptome-wide gene expression along with T cell function *in vitro* and *in vivo* will further elucidate the benefits of *μVS* and *eμVS* relative to electroporation.

## Methods

### Splitter plate device design & simulation

A total of five splitter plate device CAD geometries were designed and constructed using OnShape software. The geometries were computationally meshed and simulated with OpenFOAM software to evaluate and investigate the influence of vortex shedding on cell viability, mRNA delivery and subsequent eGFP expression. Splitter ratios for these devices were 1.0, 0.75, 0.5, 0.25 and 0 while separation ratios were 0, 0.75, 2, 3.25 and 10, respectively.

Our CFD computational domain consisted of structured (hexagonal) mesh with a total of 30 million grid points. Mesh independent studies were performed to ensure the numerical flow results were sufficiently resolved without the influence of mesh resolution. The finest resolutions used in the device geometry, located near the cylindrical posts, were at a 2 μm. This resolution was considered sufficiently to resolve flow shear layers with a total of four grid points across the device in the spanwise direction. The coarsest resolutions, located near the inlet and outlet channels of the device, were set at 8 μm. Slip boundary conditions were not used on walls except for the inlet and outlet channels, which had pressures of 134.7 and 14.7 PSIA, respectively, to create a 120 PSIG pressure drop in each device. A single-phase laminar Reynolds number was used with Opti-MEM at 23°C (ρ = 1011.4 kg m^-3^, *μ* = 9.586 x 10^4^ kg m^-1^ s^-1^) as processing fluid. 3D transient simulations were performed with OpenFOAM 5.0’s transient PimpleFOAM single phase solver. A total of 3.5 ms flow through time was simulated with a time-step of 1 μs. An initial transient time of 1.5ms was necessary to allow flow to become fully developed with the numerical schemes. Therefore, a total of 2.0 ms of statistical numerical data was collected to resolve flow structures behaviours with a flow spectral frequency from 5 kHz to 500 kHz. Numerical solutions required 19,800 to 22,800 CPU-hrs over 4 days with 198 parallel cores on Rescale and Amazon Web Services supercomputers. Rescale’s C4 Instant cluster type was used with Intel Xeon 2.9Hz CPUs with 3.8 GB of memory per core.

Vortex shedding fluctuations were quantified based on hydrodynamics fluctuating forces acting on cylindrical posts. Vortex structure scales were visualized qualitatively with the aid of spanwise vorticity fluctuation contours and Q-criterion iso-surfaces. To quantify the hydrodynamics performance of each splitter plate device, a performance indicator referred to as the Post Near Wake Indicator (PNWI) was created (SI - Detailed Simulation Analysis). PNWI values were calculated for each splitter plate device and a ratio was created relative to a baseline configuration (splitter plate ratio = 1.0) to rank their hydrodynamic performance.

### *μVS* device fabrication

Devices were fabricated using previously reported, industry standard semiconductor techniques^4^. Briefly, *μVS* device and device features (posts, channel thickness, inlet and outlet channels) were manufactured by (1) generated a digital rendering of the microfluidic using Onshape CAD software, (2) preparing a wafer substrate and (3) performing a series of metallisation, inspection, photolithography, liftoff, and laser drilling on the substrate.

### eGFP mRNA & *μVS* splitter plate study

Primary human CD3+ T cells were isolated from donor PBMCs via immunomagnetic negative selection (Stem Cell Tech). For thaw and culture, cryopreserved T cells were expanded using anti-CD3/CD28 dynabead T cell activator (ThermoFisher) with standard conditions for 2 days, followed by debeading and *μVS*.

All solutions processed through the device and device were filtered prior to use with a 0.22 μm filter to remove particulates. For device processing, T cells were debeaded, washed, resuspended in Opti-MEM (Thermofisher, 31985062) and filtered with a sterile 40 μm cell strainer. eGFP-encoding mRNA (TriLink) was delivered at 200 μg mL^-1^(50 μg mRNA) into activated T cells at 1.5 x 10^7^ cells mL^-1^ (3.75 x 10^6^ cells) in a total volume of 250 μL with a driving pressure of 120 PSIA. Sample rig and tubing were sterilized before use via 70% ethanol wipe down and flush. Immediately prior to *μVS*, samples were mixed thoroughly by gentle pipetting, mounted in the sample reservoirs and driven pneumatically though the device for intracellular delivery by *μVS*. Processed samples were collected in warmed (37°C) media (XVIV020 + 5% human serum, Lonza), and cultured at 1.0 x 10^6^ cells mL^-1^ supplemented with IL-2 at 100IU mL^-1^(Peprotech). Each splitter plate condition was evaluated in triplicate. eGFP and cell viability levels were quantified by propidium iodide viability staining (ThermoFisher) and flow cytometry (Attune, ThermoFisher Scientific) at 24 hours post-*μVS*.

### Genome editing with CRISPR Cas9, *μVS* and electroporation

Cas9 (Invitrogen) and TRAC-1-specific single-guide RNAs (sgRNAs, Synthego), as a Cas9-RNP complex (1:1 Cas9:gRNA molar ratio) were delivered primary human CD3+ T cells after 2 days of Anti-CD3/-CD28 Dynabead (ThermoFisher) stimulation. Dynabeads were removed by placing cells on a cell separation magnet for 2-5 mins. Immediately before *μVS*. de-beaded cells were centrifuged for 10 min at 1500rpm, aspirated, resuspended in Opti-MEM and filtered with a 40μm cell strainer. Cas9-RNPs were delivered at 5.0 x 10^-5^ pmols RNP cell^-1^ to 5.0 x 10^6^ activated T cells sample^-1^ at 5.0 x 10^7^ cells mL^-1^ in a total volume of 100 μL, with an applied driving pressure of 120 PSIA. For electroporation, de-beaded cells were aspirated and resuspended in 100 μL of Lonza electroporation buffer P3. 5.0 x 10^6^ activated T cells were electroporated per cuvette using a Lonza 4D X-unit electroporation system with pulse code EH115. Processed samples were collected in warmed (37°C) media (XVIV020 + 5% human serum, Lonza), and cultured at 1.0 x 10^6^ cells mL^-1^ supplemented with IL-2 at 100IU mL^-1^(Peprotech). IL-2 at 100IU mL^-1^ was supplemented every 2 ~ 3 days after transfection. Each condition was evaluated in triplicate. Levels of TCR, CD3, PD-1 and CD25 expression and cell viability and cell expansion were quantified by live/dead staining and the following antibodies via flow cytometry: anti-TCRa/b-PE (306708, BioLegend), anti-CD3-AF700 (11-0039-42, ThermoFisher), Sytox Blue Viability Stain (S34857, ThermoFisher), anti-PD-1-APC (ThermoFisher, S34857), anti-CD25-SuperBright 702 (ThermoFisher, 67-0259-42). Flow cytometry analysis was performed on days 1, 4, 7, 10 and 14 post-transfection. Cell culture supernatant levels of IFNg were quantified using Human IFN-gamma Quantikine ELISA Kit and manufacturer’s recommendations (DIF50, R&D Systems).

### Flow cytometry

Flow cytometric analysis was performed on an Attune NxT Acoustic Focusing Cytometer (ThermoFisher). Surface staining for flow cytometry was performed by pelleting cells and resuspending in flow buffer (1% human serum in PBS) with antibodies for 30 min at 4°C in the dark. Cells were washed once in flow buffer prior to resuspension.

### *eμVS* device fabrication & operation

*eμVS* devices were designed, fabricated and operated in a similar manner without a splitter plate (Figure 4). The addition of an electric field was achieved by adding platinum interdigitated electrodes to the lid of a *μVS* flow cell design with industry standard lithography methods^4^. Due to the additional fabrication requirements, the flow cell was etched in silicon then anodically bonded to a borofloat lid (see Figure 4). Fluidic access to the flow cell was achieved by laser drilling 800 μm diameter through holes in the borofloat lid and thru silicon vias enabled electrical access. 100 nm thick and 25 μm wide platinum electrodes were spaced 125 μm apart to achieve a ratio of the electrode spacing to the flow cell height suitable to create uniform electric fields between the interdigitated electrodes^8^. The electric field was applied using a standard 120V DC power supply (Nice-Power, R-SPS1203D) coupled with a customized harness to distribute the current evenly across the platinum electrodes. In addition, we used an oscilloscope (Tektronix, TBS 1052B) connected to the leads from the DC power supply to monitor the applied electric field strength. Using this circuit we applied up to 30 V of continuous DC voltage to the electrodes with a gap space of 125 μm resulting in an maximum applied electric field strength of 2.4 kV cm^-1^.

## Acknowledgements

This work was funded in part by IndieBio (indiebio.co), SOSV (sosv.com), Jobs for NSW (jobsfornsw.com.au), Y Combinator (ycombinator.com), AusIndustry (business.gov.au), Social+Capital (socialcapital.com), Main Sequence Ventures (mseq.vc), Founders Fund’s FF Science (foundersfund.com), Axial (axialsprawl.com), MTP Connect Biomedtech Horizons 1.0 (mtpconnect.org.au/biomedtechhorizons), NSW Health Medical Device Fund (medicalresearch.nsw.gov.au/medical-devices-fund) and the National Cancer Institute (75N91020C00030) along with an array of angel investors.

The device fabrication work was performed in part at the South Australia node of the Australian National Fabrication Facility, a company established under the National Collaborative Research Infrastructure Strategy to provide nano- and micro-fabrication facilities for Australia’s researchers. The design, development and prototyping of the *μVS* and *eμVS* systems were performed by Design+Industry (design-industry.com.au) and Genesys (genesysdesign.com.au).

The authors are thankful for the simulation and computational support provided by Rescale, Amazon Web Services (AWS) and the OpenFOAM community. The authors would also like to acknowledge Geoff Facer, Hamish Hawthorn, Ben Wright, David Gottlieb, Kenneth Micklethwaite, Heidi Hagen, Dino DiCarlo and Everett Meyer for their guidance, advice and support when pursuing science in the startup environment.

The content is solely the responsibility of the authors and does not necessarily represent the views of anyone acknowledged in this section.

## Author Contributions

J.A.J., B.J.S., L.H.W., G.T.S.K. and R.S.P conceived and designed the experiments. J.A.J., B.J.S., L.H.W., F.L.P., K.H.W.J.L., G.T.S.K., and A.A.L. performed the experiments. J.A.J., B.J.S., L.H.W., F.L.P., K.H.W.J.L., G.T.S.K., A.A.L., and R.S.P analyzed the data. J.A.J., F.L.P, G.T.S.K. and R.S.P contributed materials and analysis tools. J.A.J., B.J.S., L.H.W., F.L.P, and R.S.P wrote the paper.

## Tables

*Jarrell JA et al.* Genome editing human primary T cells with microfluidic vortex shedding & CRISPR Cas9

## Supplemental Figures

**Supplemental Figure 1.**
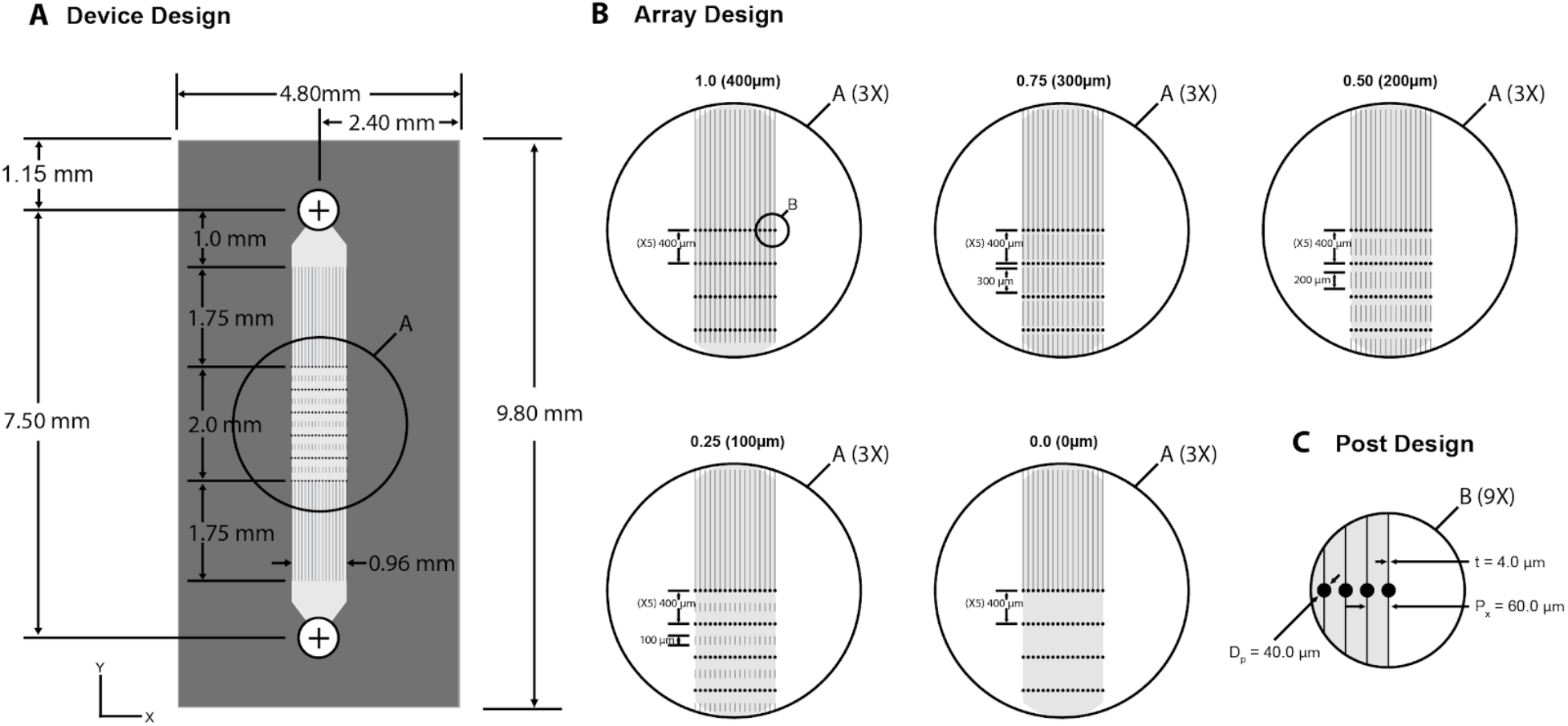
*μVS* device, array and post designs. **A,** *μVS* devices were fabricated with a 4.8 x 9.8 mm footprint with a 7.5 mm long and 40 μm deep previously-reported flow cell design^4^. Flow cells contained inlet and outlet channels flanking 6 columns of posts of 17 posts per column representing the post array region. **B**, Array designs for five *μVS* devices with varied lengths splitter plates (400-100 μm) or no splitter plate (0.0, 0 μm). **C**, All posts were identical with 40 μm diameter and depth. Splitter plate walls were of 4 μm width creating 60 μm wide channels.

**Supplemental Figure 2.**
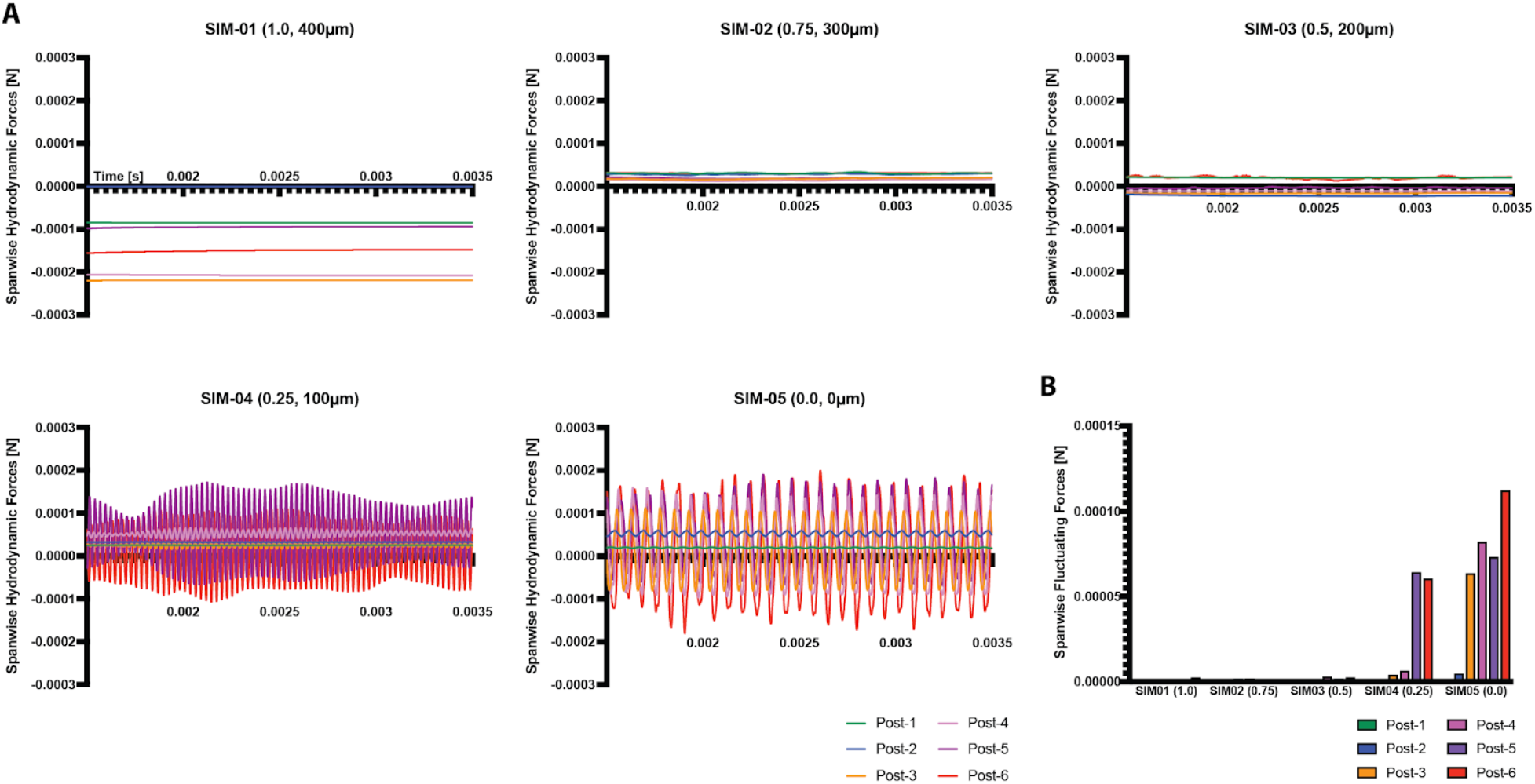
Transient force analysis of *μVS* splitter devices. (A) Spanwise hydrodynamics forces [N] on each post column of *μVS* devices with 0.0 - 1.0 splitter plate ratios as a function of time. (B) Spanwise hydrodynamic fluctuations on each post column (1 - 6) for each splitter device design.

**Supplemental Figure 3.**
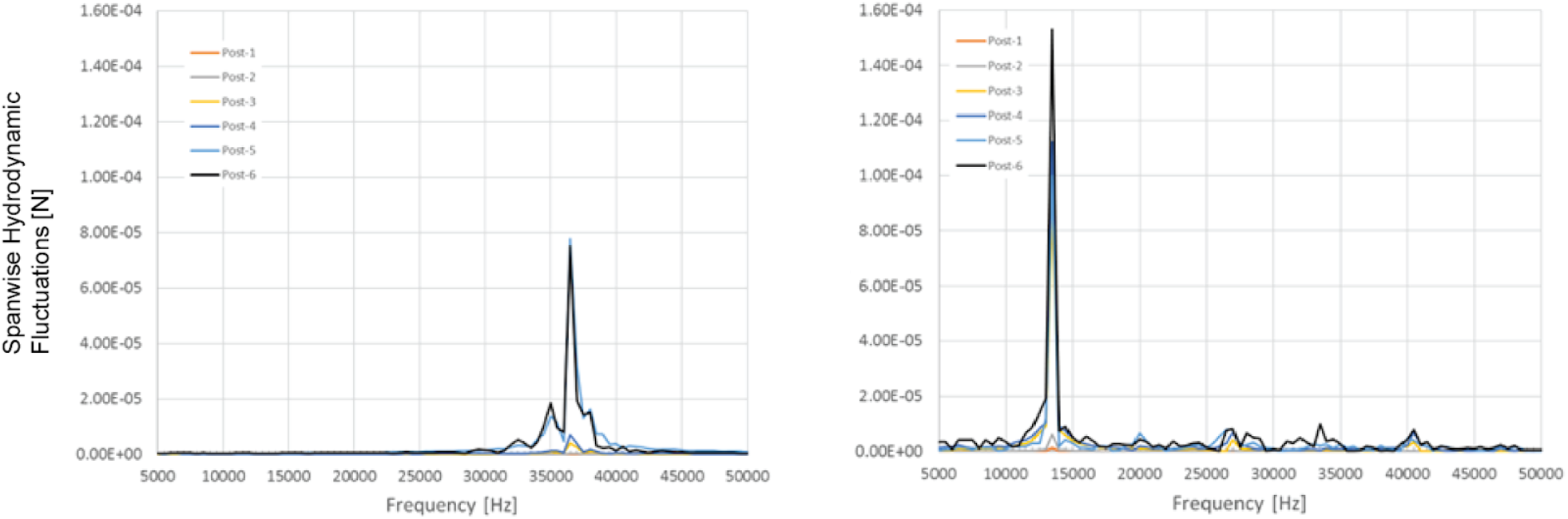
Simulated Spanwise Hydrodynamic Forces Spectral Frequency of *μVS* devices. Spanwise hydrodynamics forces spectral amplitude of both SIM04 (left, 0.25 splitter ratio) and SIM05 (right, 0.0 spitter ratio).

**Supplemental Figure 4.**
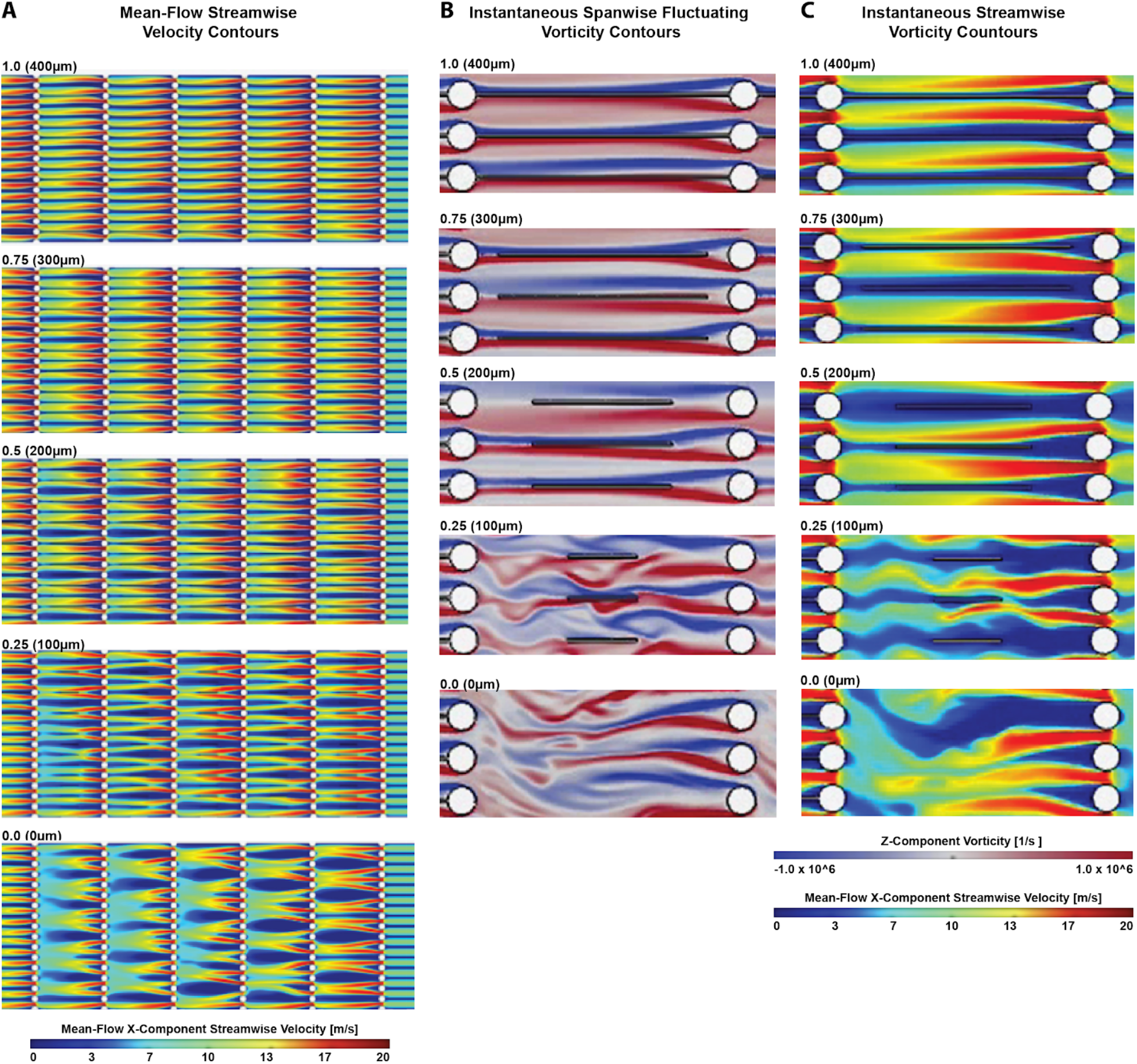
Simulated Flow-Field Contours as Mean-Flow Streamwise, Instantaneous Spanwise Fluctuating and Instantaneous Streamwise Velocity Contours of *μVS* devices. Mean-flow streamwise **A**, and instantaneous spanwise **B**, and instantaneous streamwise vorticity **C**, contour distribution for all *μVS* device designs.

**Supplemental Figure 5.**
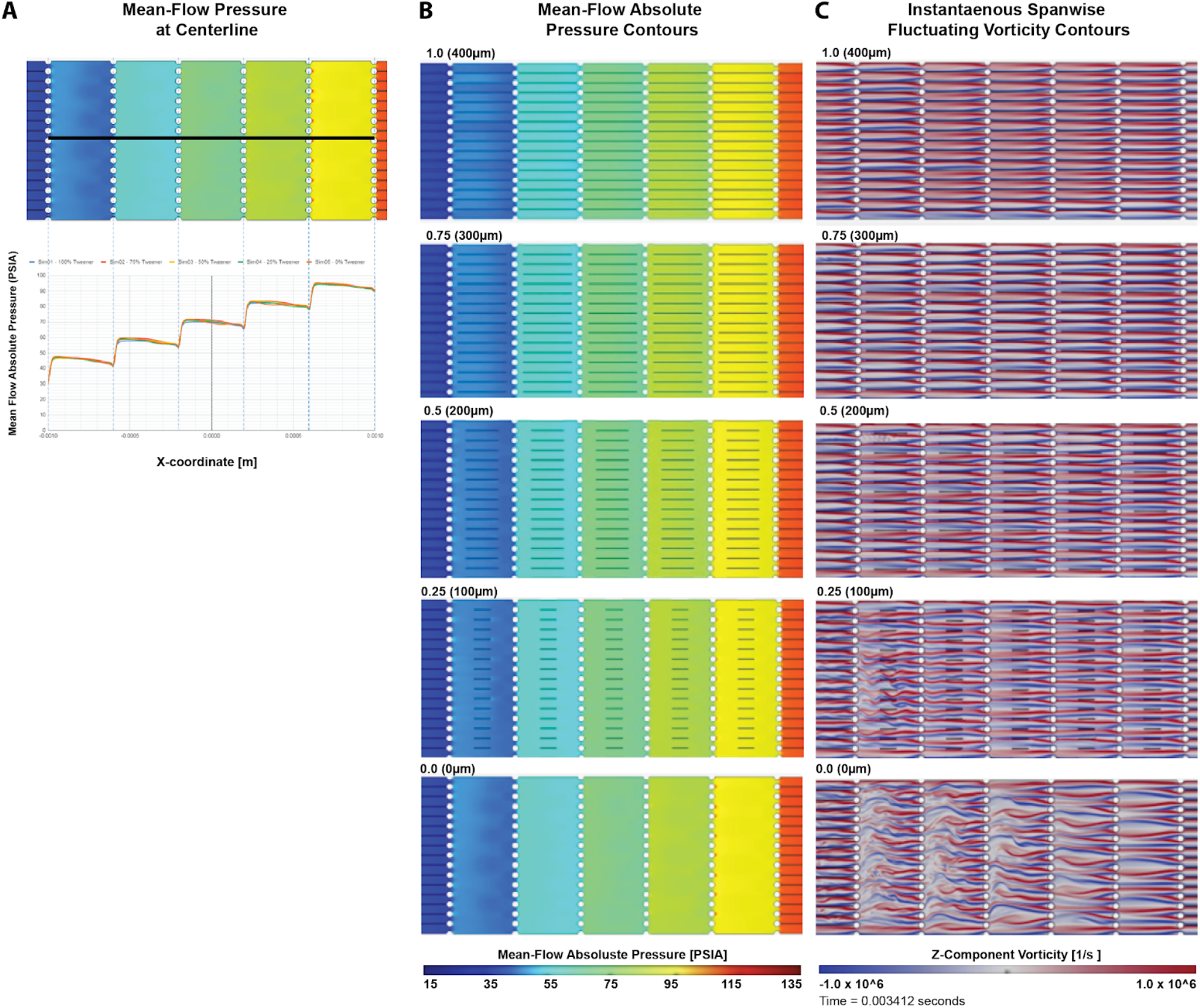
Simulated Mean-Flow Hydrodynamic Pressure Distributions and Contours and Instantaneous Spanwise Fluctuating Vorticity Contour of *μVS* devices. **A**, Mean-flow hydrodynamics pressure distribution at the centerline of the *μVS* device. Distribution is from the 1st to 6th (of 6) post-columns. Hydrodynamic pressure uniformly distributed in spanwise direction (top). Local pressure drops were detected across each post column in the streamwise direction for all design configurations (bottom). **B**, Absolute pressure and **C**, instantaneous spanwise fluctuating vorticity magnitudes in the flow fields pseudo-colored for all *μVS* devices designs.

**Supplemental Figure 6.**
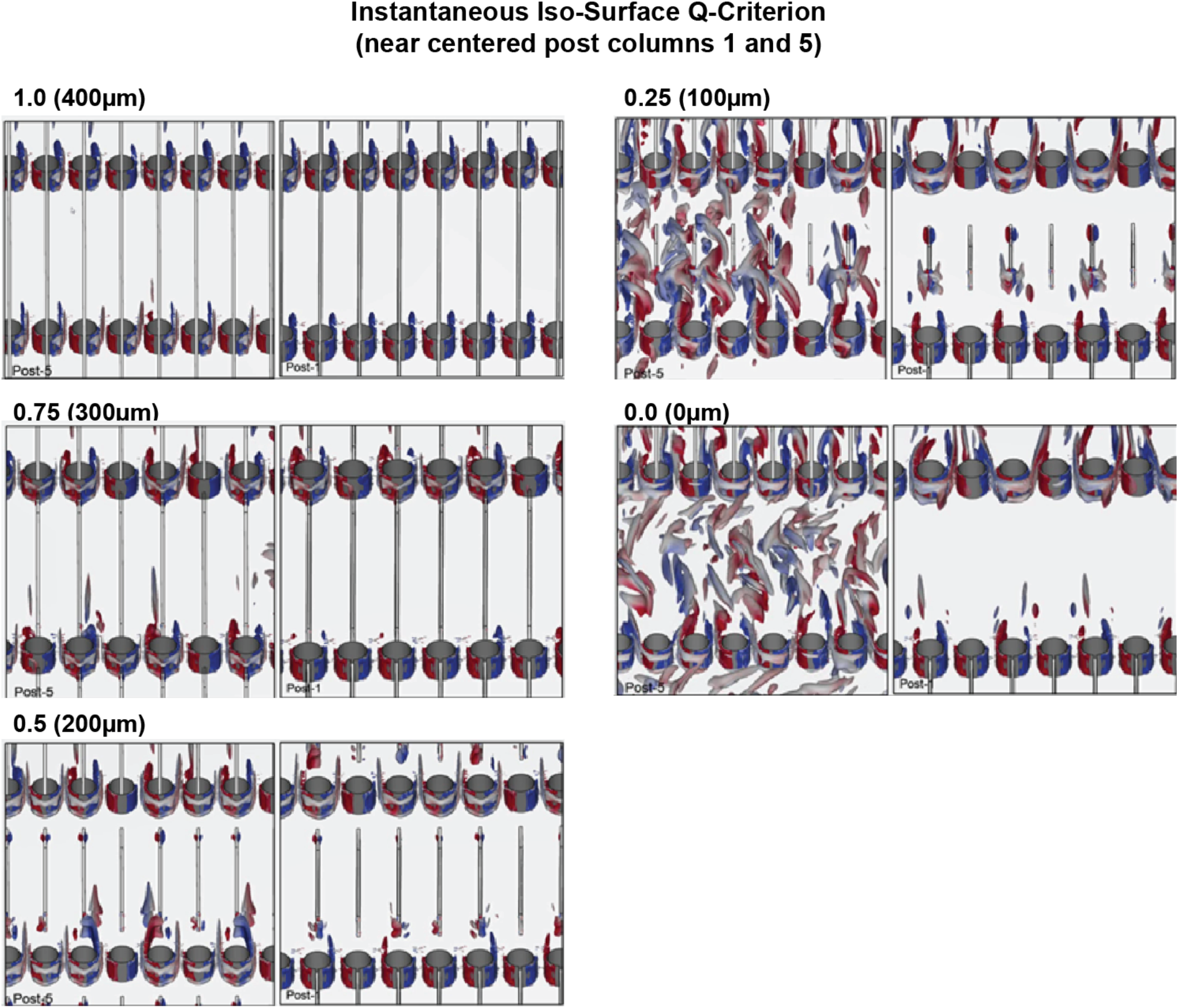
Simulated Q-criterion Iso-surfaces to visualize vortex structures in *μVS* devices. Q-criterion in flow-field simulated to visualize structures in three-dimensional space. Simulation images show Q-criterion iso-surface at post-columns 1 (right) and 5 (left) for all *μVS* device designs.

## Supplemental Methods

### Supplemental Method 1. Detailed Simulation Analysis

#### Centerline Pressure Distribution

Supplemental Figure 1 shows the mean-flow hydrodynamics pressure distribution at the centerline of the device. The distribution depicted the pressure variations at the first to last column. The presence of the splitter plates had minimal impact on the hydrodynamics pressure distribution. Local pressure drops were consistently identified across each post column in the streamwise direction for all device designs. Generally speaking, such local pressure drops varied at ~20 psig (1.4 ATM) at by-pass flow duration of 2 μs for all device designs. The local pressure drop was largely dictated by the local flow restriction due to closely aligned posts positioned along each column.

In between columns, the hydrodynamics pressures were uniformly distributed in the spanwise direction. Interestingly, these distributions remained uniform in the absence and presence of vortex shedding. Along the centerline of the device, local pressure recovery developed in the streamwise direction extending downstream into the far-field region starting from the base of each post. Such pressure recovery development was located at the far-field wake formation as flow convected downstream.

#### Mean-flow Hydrodynamics Pressure Contour and Instantaneous Spanwise Fluctuating Vorticity Contours

Supplemental Figure 1 shows flow-fields colored by absolute pressure and instantaneous spanwise fluctuating vorticity magnitude for all devices. Hydrodynamics pressure in between columns were uniformly distributed in spanwise direction for all configurations in the absence and presence of vortex shedding conditions.

As indicated in Supplemental Figure 1, fluctuating vortices were predominant in both 0.0 and 0.25 splitter ratio devices. Such spanwise fluctuations were considered the footprint of vortex shedding pairs in their oscillating modes. Red and blue colors in the contours represented counter-rotating vortices pairs with the same strength magnitude. Conversely, hydrodynamics vorticities remain undistorted (remain stationary without oscillation) in the 0.50, 0.75 and 1.0 splitter ratio devices in spanwise direction behind a majority of posts. In particular, more dominant vorticity fluctuations at the last 3 post columns were present in the 0.25 splitter ratio devices whereas uniform fluctuation behaviors across all columns was observed in the 0.5 splitter ratio device.

Therefore, we can conclude that in the presence of the vortex shedding (i.e. 0.25 and 0.0 splitter ratio), the uniform hydrodynamics pressure distribution is undisturbed since the vortex structures are fluctuating in equal and opposite strength and direction at both symmetrical sides of the post. It is expected that such a synchronous oscillating pattern of the vortex shedding in both 0.0 and 0.25 splitter cases shall not disrupt the uniformity of the pressure distribution that is similarly found in 0.5-1.0 splitter ratio cases.

#### Mean-flow Streamwise Velocity Contours

Supplemental Figure 2 shows the mean-flow streamwise velocity distributions for all design configurations. The mean-flow streamwise velocity contours provide insight to the extent of wake formation regions in the device. In the 0% splitter ratio case, the wake regions were widely spreaded in both far and near-field regions indicated by low negative streamwise velocity colored by blue. In contrast, both the 1.0 and 0.75 splitter ratio devices had fewer and narrower wake regions in most fluid domains compared to all other devices. In general, the number of wide wakes found in the domain increased with the reduction in splitter ratio. The presence of a splitter plate at the finite length of 0.50-1.0 splitter ratios showed an attenuation of near-field vortex shedding and therefore prohibited downstream (far-field) wake formation.

The large number of wide wake regions present in the 0.0 splitter ratio device is speculated to permit and promote the *μVS* intracellular delivery process. It is anticipated that the vortex shedding observed inside the wide wake regions are likely to prolong the flow duration (increase cell residence duration) for the intracellular delivery process to occur. Therefore, it is critical to conduct a set of studies with different column pitch separations for achieving the optimal performance. This work is currently ongoing.

Instantaneous flow parameters such as vorticity and streamwise velocity are useful to evaluate vortex behaviours in both near-field and far-field regions. Supplemental Figure 2 shows a close up view of a pair of spanwise oscillating vortex structures identified in 0.0 and 0.25 splitter ratio devices. In the 0.0 splitter ratio device, the spanwise oscillating vortex pairs are present, starting from each side of the post. Such vortex shedding oscillation behavior extended further downstream to the free wake region. Far-field wake development also occurred in devices with separation ratios >3.2. In all other cases, the near-field vortex shedding was attenuated in near-field and prohibited any further wake development in far-field.

#### Q-criterion Iso-Surfaces

Q-criterion detection technique was used to visualize vortex structures in three-dimensional space. Supplemental Figure 3 demonstrates the 3D vortex structures detected in between columns 1 and 2 (right) and 5 and 6 (left). Large scale vortices were detected between column 5 and 6 for 0.25 and 0.0 splitter ratio devices. Large scale vortices were found to behave in coherence as expected. In contrast, minimal vortex structures were detected upstream between column 1 and 2. These Q-criterion results were consistent with the spanwise vorticity contours previously observed and described above. For 0.25 and 0.0 splitter ratio devices, coherent vortices oscillated at much higher amplitude at downstream columns than in upstream columns.

### Supplemental Method 2. Post Near Wake Indicator (PNWI)

PNWI is defined as a Post Near Wake Indicator derived from the hydrodynamics fluctuating forces. The PNWI % calculation is based on the following steps:

1. Time history of total spanwise absolute hydrodynamics forces for each post column is recorded.
2. The standard deviation of total spanwise absolute hydrodynamics forces for each column is computed. This result gives hydrodynamics fluctuations about the mean.
3. Ratio to 0% tweener for each case is computed based on the standard deviation computed from Step 2.
4. This ratio is named “Post Near-Wake Indicator (PNWI)”. The general formula of PNWI is: ^*PNWI*% = 00% √∑(*F-F*)^

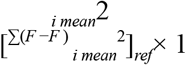

where *F* is absolute hydrodynamics forces measured at each post and is averaged (mean) *_i_ F_mean_* hydrodynamics forces measured at each post.

## References

1. Stewart, Martin P., et al. ‘ “*In vitro* and *ex vivo* strategies for intracellular delivery.” Nature 538.7624 (2016): 183–192.

2. Ghassemi, Saba, et al. “Reducing ex vivo culture improves the antileukemic activity of chimeric antigen receptor (CAR) T cells.” Cancer immunology research 6.9 (2018): 1100–1109.

3. Stadtmauer, Edward A., et al. “CRISPR-engineered T cells in patients with refractory cancer.” Science (2020).

4. Jarrell, Justin A., et al. “Intracellular delivery of mRNA to human primary T cells with microfluidic vortex shedding.” Scientific Reports 9.1 (2019): 1–11.

5. Unal, M. F., and D. Rockwell. “On vortex formation from a cylinder. Part 2. Control by splitter-plate interference.” Journal of Fluid Mechanics 190 (1988): 513–529.

6. DiTommaso, Tia, et al. “Cell engineering with microfluidic squeezing preserves functionality of primary immune cells in vivo.” Proceedings of the National Academy of Sciences 115.46 (2018): E10907–E10914.

7. Ding, Xiaoyun, et al. “High-throughput nuclear delivery and rapid expression of DNA via mechanical and electrical cell-membrane disruption.” Nature biomedical engineering 1.3 (2017): 1–7.

8. Adamo, Andrea, et al. “Flow-through comb electroporation device for delivery of macromolecules.” Analytical chemistry 85.3 (2013): 1637–1641.

9. Barrangou, Rodolphe, and Jennifer A. Doudna. “Applications of CRISPR technologies in research and beyond.” Nature biotechnology 34.9 (2016): 933–941.

10. Roth, Theodore L., et al. “Reprogramming human T cell function and specificity with non-viral genome targeting.” Nature 559.7714 (2018): 405–409.

11. Zhang, Mingce, et al. “The impact of Nucleofection® on the activation state of primary human CD4 T cells.” Journal of .mmunological methods 408 (2014): 123–131.

12. Riemen, Gudula, et al. “Buffer solution for electroporation and a method comprising the use of the same.” U.S. Patent No. 8,039,259. 18 Oct. 2011.

13. Venslauskas, Mindaugas S., and Saulius Šatkauskas. “Mechanisms of transfer of bioactive molecules through the cell membrane by electroporation.” European Biophysics Journal 44.5 (2015): 277–289.

14. Lissandrello, Charles A., et al. “High-throughput continuous-flow microfluidic electroporation of mRNA into primary human T cells for applications in cellular therapy manufacturing.” Scientific reports 10.1 (2020): 1–16.

15. Sutrave, G., et al. “Piggybat transposase for the generation of CD19 specific chimeric antigen receptor T cells.” Cytotherapy 21.5 (2019): S17.

